# Molecular mechanism of the anti-lung cancer effect of Jin Ning Fang based on network pharmacology and experimental verification

**DOI:** 10.1101/2021.07.26.453881

**Authors:** Chunxiao Wu, Qiquan Yu, Weizhen Shou, Kun Zhang, Yang Li, Wentao Guo, Qi Bao

## Abstract

**Background:** Jin Ning Fang (JNF) is widely used as an adjuvant therapy for lung cancer. However, its molecular mechanism against lung cancer is still unclear.

**Methods:** The chemical compounds JNF were screened from the TCMSP database and its target proteins were then predicted. The genes related to lung cancer were collected from the CTD and DisGeNET databases. Next, targets were integrated with disease-related genes to obtain candidate genes. Functional enrichment and protein-protein interaction (PPI) analysis were also performed, followed by construction of pharmacological network. Meanwhile, Autodock was used to assess the affinity between targets and compound. Finally, the anti-cancer effect of JNF on lung cancer cells was detected and some predicted key genes was validated by using real-time PCR.

**Results:** Twenty-five overlapping targets were obtained, and pathway analysis showed that JNF might exert its anti-cancer function by regulating some biological pathways, such as apoptosis pathway. PPI and pharmacological network revealed several core targets (such as AKT1, AR, and ESR1) and three compounds (quercetin, calcium carbonate, and beta-sitosterol). Then, beta-sitosterol had a high affinity with AKT1, AR, and ESR1. Further *in vitro* experiments confirmed that JNF could inhibit proliferation and promote apoptosis of A549 cells. The expression of FDPS, PIM1, VCAM1, SLC29A1, NQO1, and ESR1 were significantly decreased, while mRNA level of AR and ANPEP were markedly increased after JNF treatment.

**Conclusion:** JNF may exert anti-lung cancer effect through multiple targets and pathways, and identified genes may be used as potential biomarkers for diagnosis and treatment of lung cancer.

## Introduction

Lung cancer is a leading cause of cancer death in China, with significantly increased incidence and mortality rates in recent years, especially with the aggravation of air pollution (Cao and Chen 2019; Guo et al. 2019). In recent years, as the main or adjuvant medicine, traditional Chinese medicine (TCM) has been widely used in the treatment of lung cancer with the advantage of low side effects (Su et al. 2020; Zhang et al. 2018). However, owing to the complex chemical composition and diversity of TCM prescriptions, great challenges have been encountered in revealing the potential active compounds and anticancer mechanism of TCM.

The TCM prescription JNF shows the effects of clearing heat and removing toxicity. It is composed of five Chinese classic herbs, and our previous study has indicated that JNF could inhibit the proliferation of human lung cancer cell line (95-D) within 24-48 h under various concentrations. JNF also regulated the expression level of Caspase 3 and Caspase 8 to promote cell apoptosis (Wu et al. 2015). We also found that JNF combined with postoperative adjuvant chemotherapy inhibited metastasis recurrence in patients with stage III non-small cell lung cancer (NSCLC) after surgery; in addition, it indeed exerted a relatively stable and controlled effect on the main serum tumor markers, and thus could significantly improve the quality of life of patients (Shou et al. 2014). These studies further indicated the potential therapeutic effects of JNF against lung cancer. However, the molecular mechanism of the anti-lung cancer effect of JNF is still unclear.

Several methods can be employed to investigate the mechanisms of the antitumor effects of TCM, including network pharmacology. The concept of network pharmacology was first proposed by Hopkins in 2007 (Hopkins 2007). Subsequently, herb network in TCM research framework was updated as “Phenotype network,” and “Network target” was proposed as a new concept for clarifying the herb formulas of TCM (Yoo et al. 2018). Notably, network pharmacology provides support for rational drug design and a better understanding of the mechanisms of multiple drug actions. Thus, we utilized network pharmacology to explore the potential pharmacological mechanism of JNF against lung cancer. First, the active compounds of JNF were screened from the Traditional Chinese Medicine Systems Pharmacology Database and Analysis Platform (TCMSP) and the Traditional Chinese Medicine Information Database (TCMID). Next, the target proteins of the compounds were predicted and integrated with lung cancer-related genes to obtain candidate genes. Subsequently, these candidate genes were subjected to functional enrichment analysis and protein-protein interaction (PPI) analysis, followed by construction of pharmacological network. Meanwhile, molecular docking was conducted to predict the interactions between the compounds and their target proteins. Finally, in vitro experiments were conducted to observe the inhibitory effect of JNF on the growth of lung cancer cells. In addition, real-time PCR (qPCR) was performed to detect the expression level of identified genes. Our study might help clarify the potential mechanism of the anti-lung cancer effect of JNF and facilitate the development of novel drugs.

## Material and methods

### Active ingredients of JNF

The chemical compounds of JNF were obtained from TCMSP (http://183.129.215.33/tcmid/search/) (Ru et al. 2014) that is an efficient system pharmacology platform consisting of 499 Chinese herbs registered in the Chinese Pharmacopoeia, with 29,384 ingredients, 3,311 targets, and 837 related diseases. For herbs not recorded in TCMSP, we searched the TCMID (http://183.129.215.33/tcmid/search/) database for complete information. Furthermore, the active compounds of JNF were queried using absorption, distribution, metabolism, and excretion (ADME) analysis (Yamashita and Hashida 2004). In this analysis, the active compounds of JNF were selected with the criteria of oral bioavailability ≥ 30% and drug-likeness ≥ 0.18.

### Identification of target proteins

The target proteins corresponding to the chemical compounds were predicted based on the TCMSP and Bioinformatics Analysis Tool for Molecular Mechanism (BATMAN) database (http://bionet.ncpsb.org/batman-tcm/) (Liu et al. 2016). In brief, targets were obtained from the DrugBank, Kyoto Encyclopedia of Genes and Genomes (KEGG), and Therapeutic Target Database (TTD) databases, and then the predicted targets were sorted in a descending order of score, which was calculated by the similarity between the target interaction and known target. Target proteins with a score of more than 20 were selected for subsequent analysis.

### Candidate target collection

The Comparative Toxicogenomics Database (CTD, 2018 update, http://ctdbase.org/) (Davis et al. 2019) is aimed to promote the understanding of information on environmental risks affecting the human health, and it provides information on chemical-protein/gene interactions as well as chemical–disease and gene–disease relationships to help analyses of disease mechanisms associated with environmental impact. In addition, DisGeNET is a comprehensive platform that integrates human disease-related genes and mutation information. The genes related to “lung cancer” were searched in the CTD and DisGeNET databases (Piñero et al. 2015). Notably, genes were sorted according to the inference score (Barabasi and Albert 1999; Li and Liang 2009). The inference score is calculated by the log-transformed product of two public neighborhood statistics, which is used to evaluate the functional correlation among proteins in the PPI network. Next, genes with an inference score of ≥ 20 were selected and then integrated with the targets of the compounds. Overlapping genes were selected as candidate targets for further analysis.

### Functional enrichment analysis of candidate genes

To elucidate the biological function of the candidate genes, functional enrichment analysis was performed. Gene ontology (GO) function analysis was conducted using the Cytoscape plugins ClueGO and CluePedia (Bindea et al. 2009; Ashburner et al. 2000). The relevant biological process (GO-BP) was selected, and P.adjust ≤ 0.01 was set as the cut-off criteria. Owing to the complexity of the structure of the GO tree (directed acyclic graph), the global options GO levels 3-8 were selected in the study, and the GO functional network was visualized by Cytoscape. Kappa coefficient was used for consistency analysis and for measuring classification accuracy. Kappa coefficient was calculated based on the confusion matrix. In the ClueGO plugin, Kappa coefficients showed the relationship between GO terms based on overlapping genes. Higher kappa coefficients indicate stronger term association. In addition, KEGG pathway was analyzed by using clusterProfiler, with a value of P ≤ 0.05 set as the threshold.

### PPI analysis

The STRING database (version 10.0, http://www.string-db.org/) provides predicted and experimental information of protein interactions. Required confidence (combined score) of > 0.9 was selected as the threshold for PPI analysis. Next, the PPI network was visualized by Cytoscape.

### Pharmacological network construction

To better understand the molecular mechanism of JNF against lung cancer, a pharmacological network of the herb compounds, targets, and pathways was constructed using Cytoscape. Important active components of JNF decoction were revealed in the pharmacology network based on their role in the regulation of the related target proteins.

### Validation of compound-target via molecular docking

The three-dimensional (3D) structures of the candidate genes were downloaded from the RCSB Protein Data Bank database (http://www.rcsb.org/), and the molecular structures of the key compounds were downloaded from the DrugBank database (https://go.drugbank.com/). Subsequently, the core compound was used as a ligand and the protein corresponding to the core target gene was used as a receptor. Molecular docking was performed using Autodock (version 4.2.6).

### Preparation of the JNF

JNF is composed of five dried medicinal herbs including *Selaginella doederleinii* Hieron. (Selaginellaceae), 30 g; *Salvia chinensis* Benth. (Labiatae), 20 g; *Bufo bufo gargarizans* Cantor (Bufonidae), 5 g; *Bombyx batryticatus* (Bombycidae), 5 g; and *Ostrea gigas* Thunberg (Ostreidae), 10 g. These herbs were mixed and extracted with boiling water for twice. Next, the filtrates were combined and concentrated to JNF decoction (1.5 g/mL of crude drug), and then stored at 4°C.

### Acquisition of JNF-containing serum

Four adult male Sprague-Dawley (SD) rats (240-260 g) were purchased from Shanghai Keting Biological Technology Co., Ltd. and allowed to acclimate for 1 week under standard condition. Then, rats were randomly divided into two groups (*n* = 2), including control group and Jing Ning Fang group. In brief, the rats in the JNF group were given intragastric administration with JNF decoction of 14g/kg/d (equal to clinical dosage), and rats in the control group were given the same volume of normal saline. After 7 days of administration, rats were anesthetized with sodium pentobarbital (30 mg/kg), and blood was collected from the abdominal aorta. The collected blood was centrifuged at 3000 rpm/min for 15 min, and the supernatant was obtained. Next, the supernatant was inactivated at 56°C for 30 min, and then filtrated via a 0.22 μm membrane. Finally, the serum was dispensed into 10 mL bottles and preserved at −80°C for further use. All animal experiments were performed in accordance with the NIH Guide for the Care and Use of Laboratory Animals and the procedures were approved by the animal ethic committee of Longhua Hospital Affiliated to Shanghai TCM University.

### Cell culture and treatment

Human lung cancer cell line A549 was purposed from National Collection of Authenticated Cell Cultures (Shanghai, China). A549 cells were cultured in RPMI-1640 medium (Invitrogen) containing 10% fetal bovine serum (FBS), 100 µg/mL penicillin, and 100 mg/mL streptomycin. Cells were maintained in a humidified condition containing 5% CO_2_ at 37°C. Subsequently, the cells in the logarithmic growth phase were trypsinized, and centrifuged at 300 g for 3 min. After discarding the supernatant, 3 mL of fresh medium was added to resuspend the cells. Next, cells were cultured in 96-well plates with 4000 cells per well at 37°C with 5% CO_2_ for one day. Culture medium was then changed and cells were divided into two groups: control group (basic medium with 10% normal rat serum) and JNF group (basic medium with 10% rat medicated serum).

### CCK-8 assay

CCK-8 assay was performed to assess the cell viability. After incubation for 48 h, 100 μL of fresh serum-free medium containing 10% CCK-8 solution (5 mg/mL) was added to each well. After culturing in dark for 1 h, the absorbance was detected at 450 nm using a microplate reader (Infinite M100 PRO) and then calculated the cell viability.

### Apoptosis assay

The ratio of apoptotic cells was examined using an Annexin V-PE/7-AAD apoptosis kit (#88-8102-74; Ebioscience, AUT) according to the guide of manufacturer. In brief, the treated cells were washed with 1 mL of cold PBS, and then re-suspended in 100 μL 1× binding buffer. Next, 5 μL Annexin V-PE and 5 μL 7-AAD (50 ug/mL) were supplemented to cell suspension. After mixing, cells were incubated in dark at temperature (25°C) for 15 min. Finally, the cell samples were detected by FACSCalibur flow cytometry (BD Biosciences) and analyzed by using Flowjo7.6.1 software.

### qPCR assay

In order to detect the mRNA expression level of several key genes obtained in the pharmacological network, the qPCR analysis was conducted as previously described (Nolan et al. 2006). In brief, the treated cells were washed with PBS, and then 1 mL of Trizol was added to extract total RNA. The reverse transcription was performed using the ReverTra Ace® qPCR RT Master Mix kit (#FSQ-201; TOYOBO co., ltd.) and qPCR was conducted using the Power SYBR Green PCR Master Mix kit (#A25742; Thermo Fisher Scientific). The relative expression level of genes was normalized by using GADPH. The primer sequences for genes are listed in Table S1. *Statistical analysis*

All experiments were repeated three times and data were presented as the mean ± standard deviation (SD). Statistical analysis was performed using the Graphpad prism 5 (Graphpad Software, San Diego, CA). Statistical comparison was conducted using the unpaired *t* test or two-tailed test. Data with values of P < 0.05 were considered as statistically significant.

## Results

### Screening of active compounds

The number of chemical compounds identified for each ingredient of JNF was as follows: *Selaginella doederleinii* Hieron., 30; *Salvia chinensis* Benth., 16; *Bufo bufo gargarizans* Cantor, 24; *Ostrea gigas* Thunberg, 10; and *Bombyx batryticatus*, 4. After screening by using the ADME criteria, a total of 27 active compounds were identified from the five ingredients of JNF. Information on the chemical composition of each ingredient is shown in Table 1. In addition, two ingredients (Chinese sage herb and oyster) and 218 targets of seven important components were searched in the BATMAN database.

**Table 1.** Important chemical compounds in Jin Ning Fang decoction

| Ingredients | Before | After | Source |
| --- | --- | --- | --- |
| *Selaginella doederleinii* Hieron. | 30 | 1 | TCMSP |
| *Salvia chinensis* Benth. | 16 | 14 | TCMSP |
| *Bufo bufo gargarizans* Cantor | 24 | 5 | TCMID |
| *Ostrea gigas* Thunberg | 10 | 6 | TCMID |
| *Bombyx batryticatus* | 4 | 1 | TCMID |
| Total | 84 | 27 |  |

TCMSP: Medicine Systems Pharmacology Database and Analysis Platform; TCMID:
the Traditional Chinese Medicine Information Database.

### Screening and cross-validation of target proteins

A total of 3681 lung cancer-related genes were obtained from the CTD and DisGeNET databases. After integrating these genes and the targets of the active compound, 25 overlapping genes were obtained for further analysis (Figure 1).

**Figure 1.**
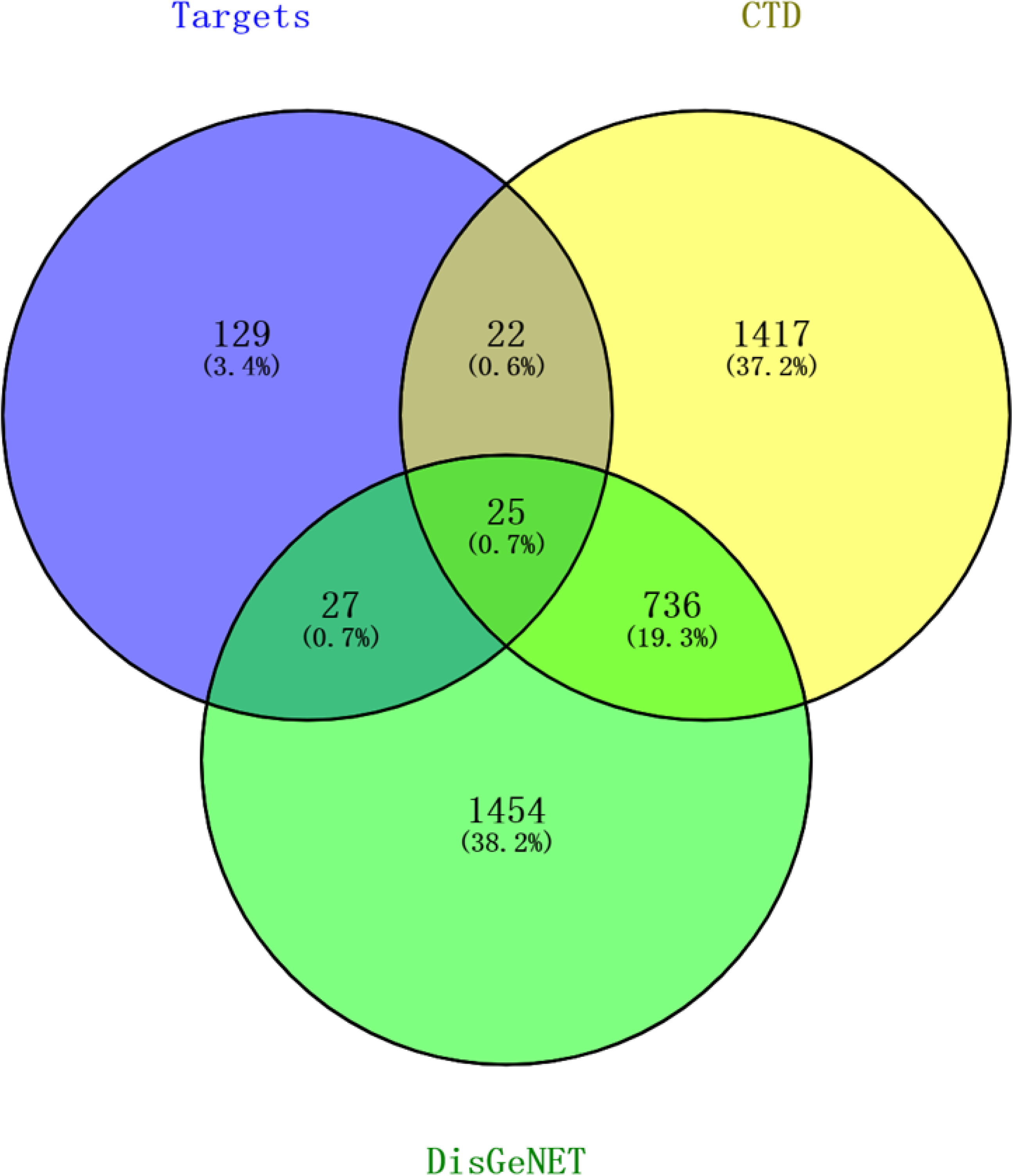
Venn diagram of predicted cross-validation targets.

### Functional enrichment pathway analysis of cross-targets

GO functional analysis revealed that these overlapping genes were significantly enriched in 45 GO functional terms (Figure 2A), and these terms were divided into seven categories based on the kappa coefficient, such as negative regulation of vitamin metabolic processes and regulation of lipid biosynthetic processes (Figure 2B). Moreover, the GO functional network showed that 19 targets were significantly enriched in 45 GO-BP terms (Figure 3A). The most enriched GO-BP terms were endogenous apoptosis signaling pathways of DNA damage, cell responses to ketones, cell responses stimulated by corticosteroids, and lymphocyte homeostasis. Pathway enrichment analysis of cross-targets was performed using clusterProfiler, and a total of 158 pathways were obtained. Pathways with P.adjust < 0.05 are presented in Figure 3B. Results showed that these genes were mainly involved in ABC transport, platinum drug resistance, and apoptosis pathways.

**Figure 2.**
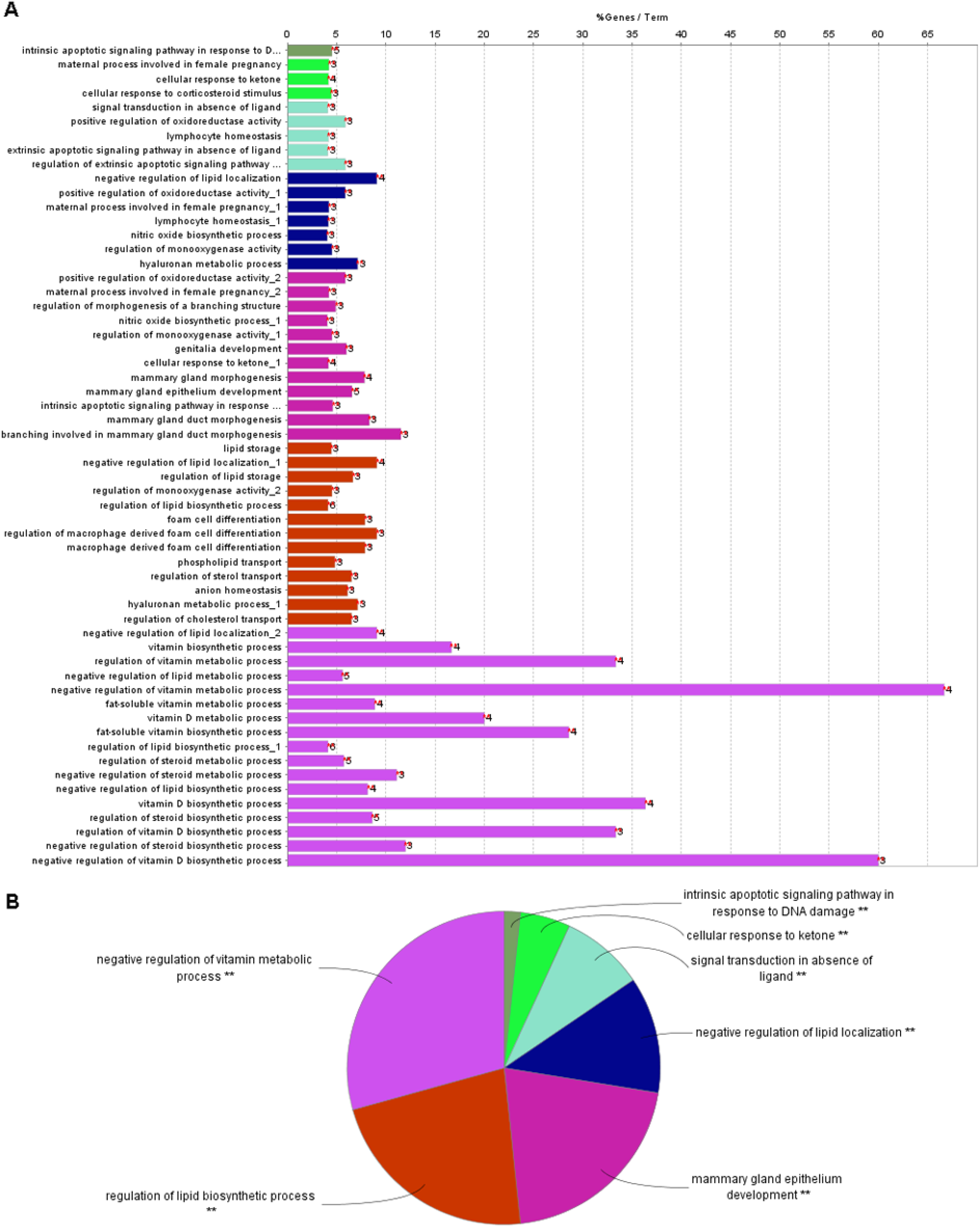
Results of Gene Ontology (GO) functional analysis. A: Histogram. The vertical axis is the GO term, and the horizontal axis is the percentage of genes enriched in each related GO functional term. GO functional grouping is distinguished by the color of the bars. The number on the right of the bar represents the number of genes enriched in each related GO functional term. * P < 0.05, ** P < 0.01. B: Pie chart.

**Figure 3.**
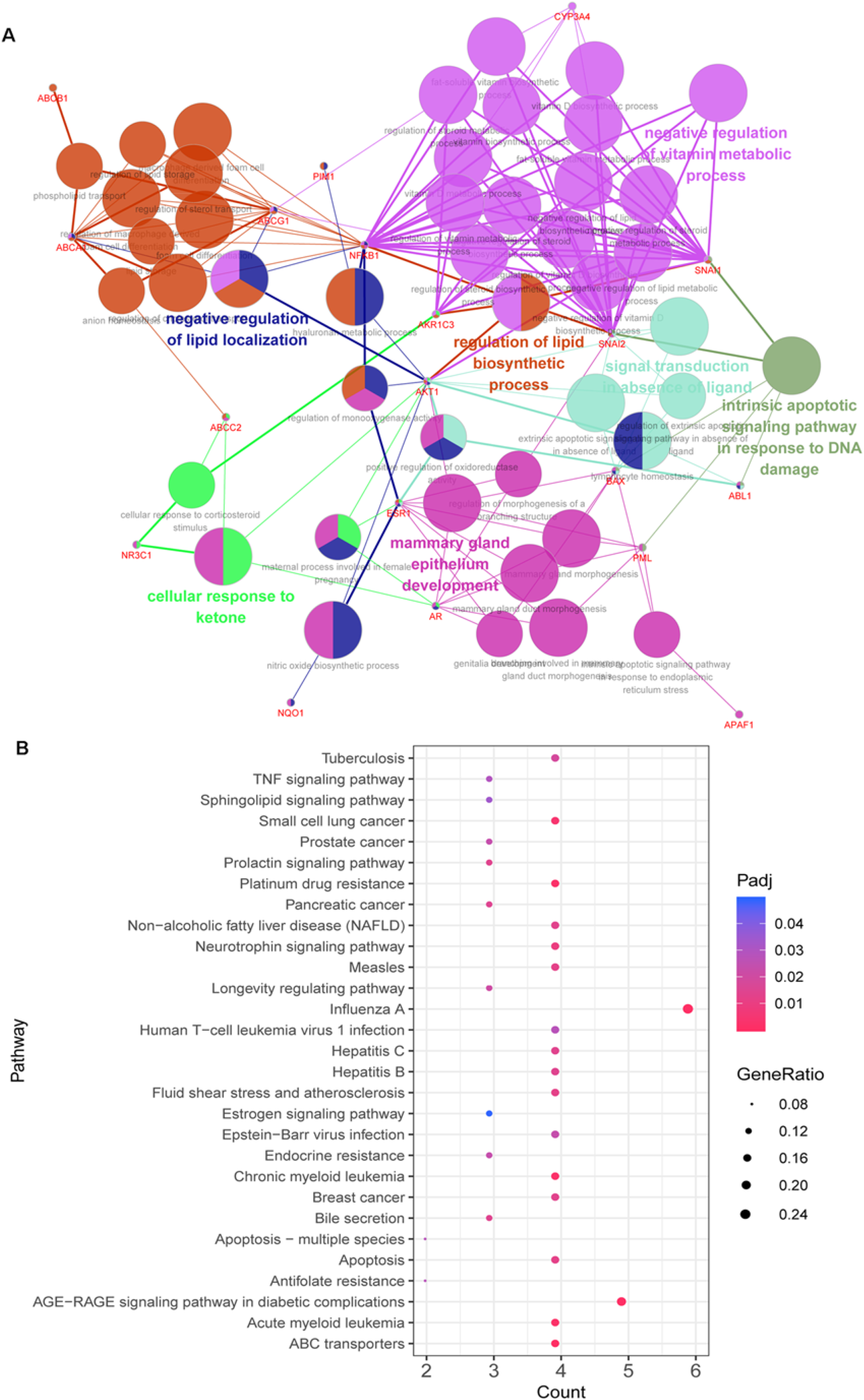
Functional enrichment analysis of 25 target proteins. A: GO-BP functional network. Larger dots indicate smaller P values. Thicker lines represent larger kappa coefficients. Diamond arrows represent regulation, and T arrows represent inhibition or negative regulation. B: KEGG pathway for target proteins. The vertical axis is the path name, and the horizontal axis is the number of enriched genes.

### PPI network analysis

Based on the 25 candidate genes, a PPI network was constructed by importing the gene ID of these targets to the STRING database. As shown in Figure 4, a total of 23 nodes and 61 interactions were obtained. The top 10 proteins with the highest degree of association are shown in Table 2, including AKT1, AR, VCAM1, and ESR1, and these proteins were recognized as hub proteins.

**Figure 4.**
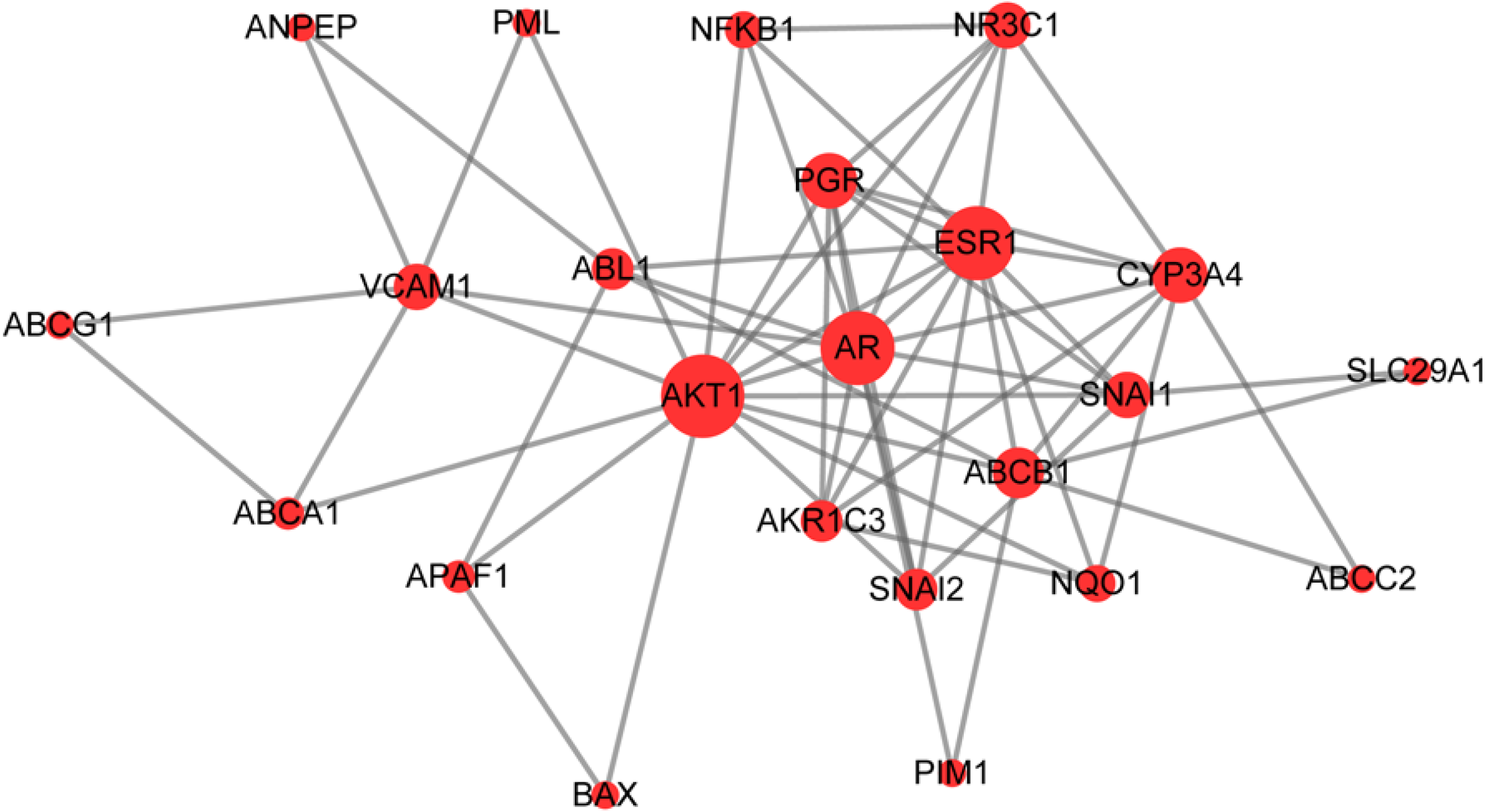
PPI network of target proteins. The size of the dots represents the degree of association with other proteins. The larger the dot, the stronger the degree of association.

**Table 2.** Top ten genes in protein-protein interaction (PPI) network

| Gene | Degree | Betweenness | Closeness |
| --- | --- | --- | --- |
| AKT1 | 14.0 | 168.17143 | 0.73333335 |
| AR | 12.0 | 81.115875 | 0.6875 |
| ESR1 | 12.0 | 42.85873 | 0.6666667 |
| CYP3A4 | 8.0 | 23.290476 | 0.52380955 |
| PGR | 8.0 | 9.179365 | 0.57894737 |
| ABCB1 | 7.0 | 56.263493 | 0.57894737 |
| VCAM1 | 6.0 | 52.404762 | 0.55 |
| SNAI1 | 6.0 | 18.542856 | 0.5365854 |
| NR3C1 | 6.0 | 3.0269842 | 0.5365854 |
| ABL1 | 5.0 | 29.566668 | 0.52380955 |

### Pharmacological network analysis

Furthermore, a pharmacological network was constructed based on ingredients, active components, target proteins, and pathways, and the results are shown in Figure 5. This network was composed of 59 nodes and 134 pairs. Among these nodes, there were 2 ingredients (Chinese sage herb and oyster), 3 chemical composition nodes (quercetin, calcium carbonate, and beta-sitosterol), 25 target proteins (including AKT1, AR, VCAM1, FDPS, and ESR1), as well as 29 pathways (including ABC transport, platinum drug resistance, and apoptosis).

**Figure 5.**
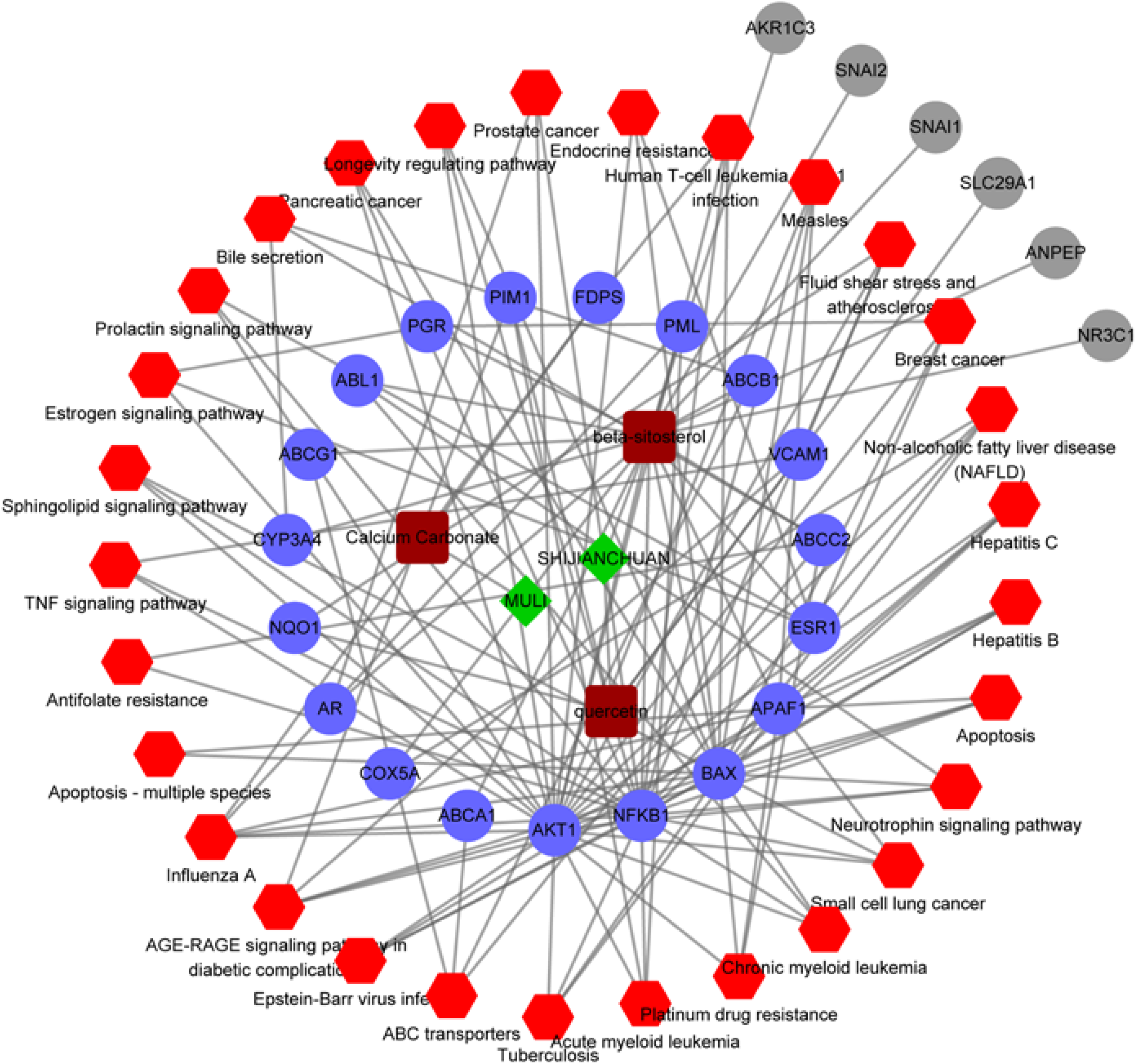
Pharmacology network. Green diamonds represent herbs, brown squares represent compounds, blue dots represent target genes, red hexagons represent KEGG pathways, and gray dots represent targets that are not involved in the pathways.

### Results of molecular docking

The above analysis revealed genes (AKT1, AR, and ESR1) and active compounds (beta-sitosterol) as key nodes. Thus, molecular docking analysis was performed using Autodock 4.2.6 to visualize the interaction between beta-sitosterol and its potential targets. A total of 10 docking models were obtained for three genes and beta-sitosterol. Binding energy was scored in the molecular docking analysis; a lower score indicates stronger binding affinity. Thus, we sorted the docking scores and selected the best docking model for the three genes (the complete data are listed in Table S2). The optimal molecular docking models are shown in Figure 6. Results showed that beta-sitosterol had a good docking with the corresponding proteins, and the key amino acids around it mainly played a role in the formation of hydrogen bonds. Hydrogen bonds minimized the energy of small compounds; thus, small compounds were the most stable. Moreover, the binding energies of AKT1, AR, and ESR1 were -10.36, -8.16, and -8.97, respectively. The docking energy scores of these compounds were relatively small, suggesting that they could stably bind to the receptor proteins and exert their effects.

**Figure 6.**
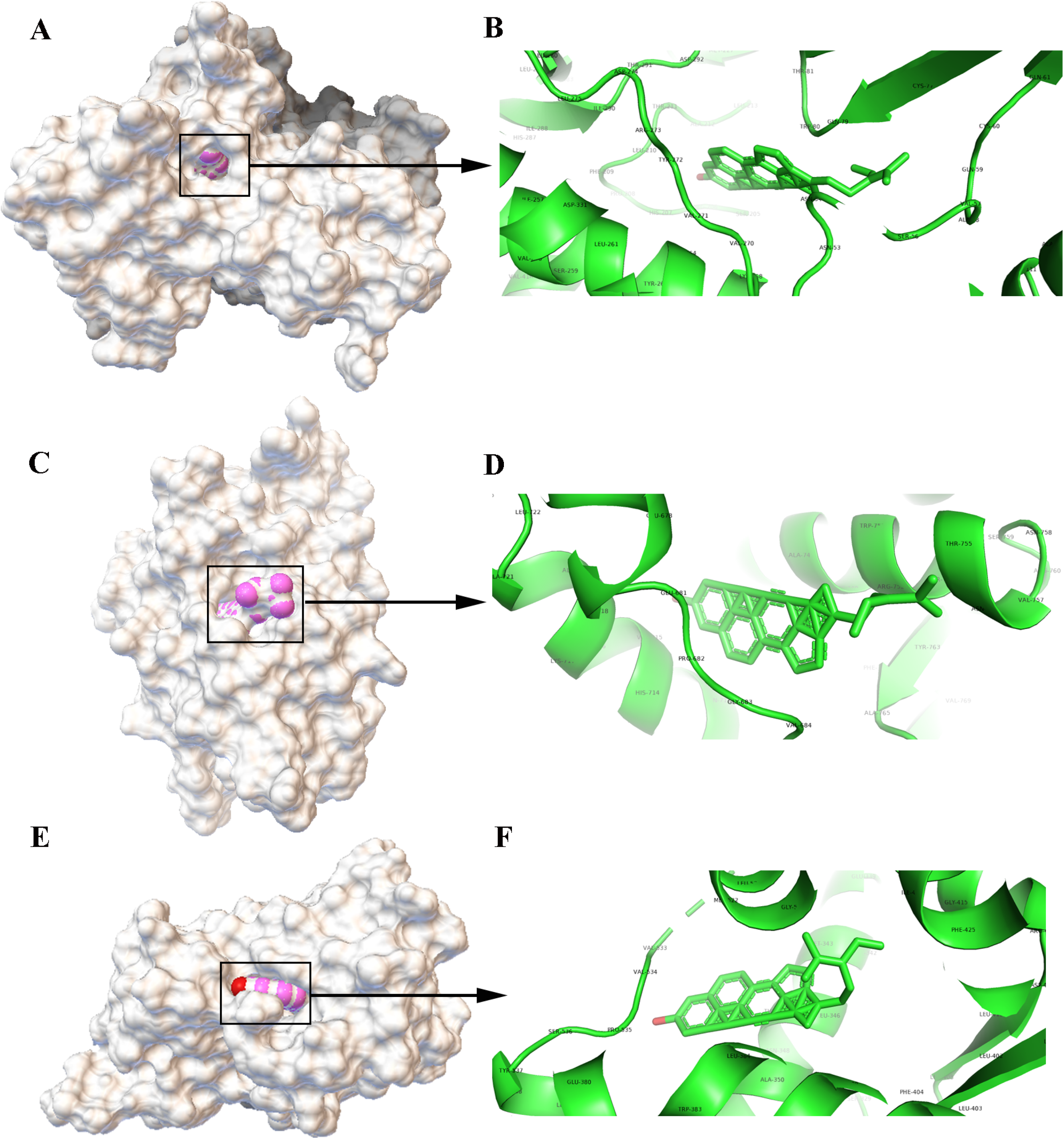
Molecular models of the binding of beta-sitosterol to its predicted protein targets. A: Global map of AKT1-beta-sitosterol molecular docking. Off-white represents the receptor, and the pink ball indicates the small ligand beta-sitosterol. B: Binding interaction between AKT1 and beta-sitosterol. C: Global map of the AR-beta-sitosterol model. D: Binding interaction between AR and beta-sitosterol. E: Global map of the ESR1-beta-sitosterol model. F: Binding interaction between ESR1 and beta-sitosterol.

### Effect of JNF on the proliferation and apoptosis of lung cancer cell line

Further, A549 cells were treated with JNF for 48 h. As shown in Figure 7A, the cell viability in JNF was significantly lower than in control group (100.00 ± 1.47 vs. 82.59 ± 2.81, P < 0.01), suggesting that JNF observably inhibited the proliferation of lung cancer cells. In addition, JNF increased the apoptosis rate in A549 cells compared with the control group (0.74 ± 0.03 vs. 2.20 ± 0.14, P < 0.01), indicating that lung cancer cells treated with JNF may induce apoptosis (Figure 7B and 7C).

**Figure 7.**
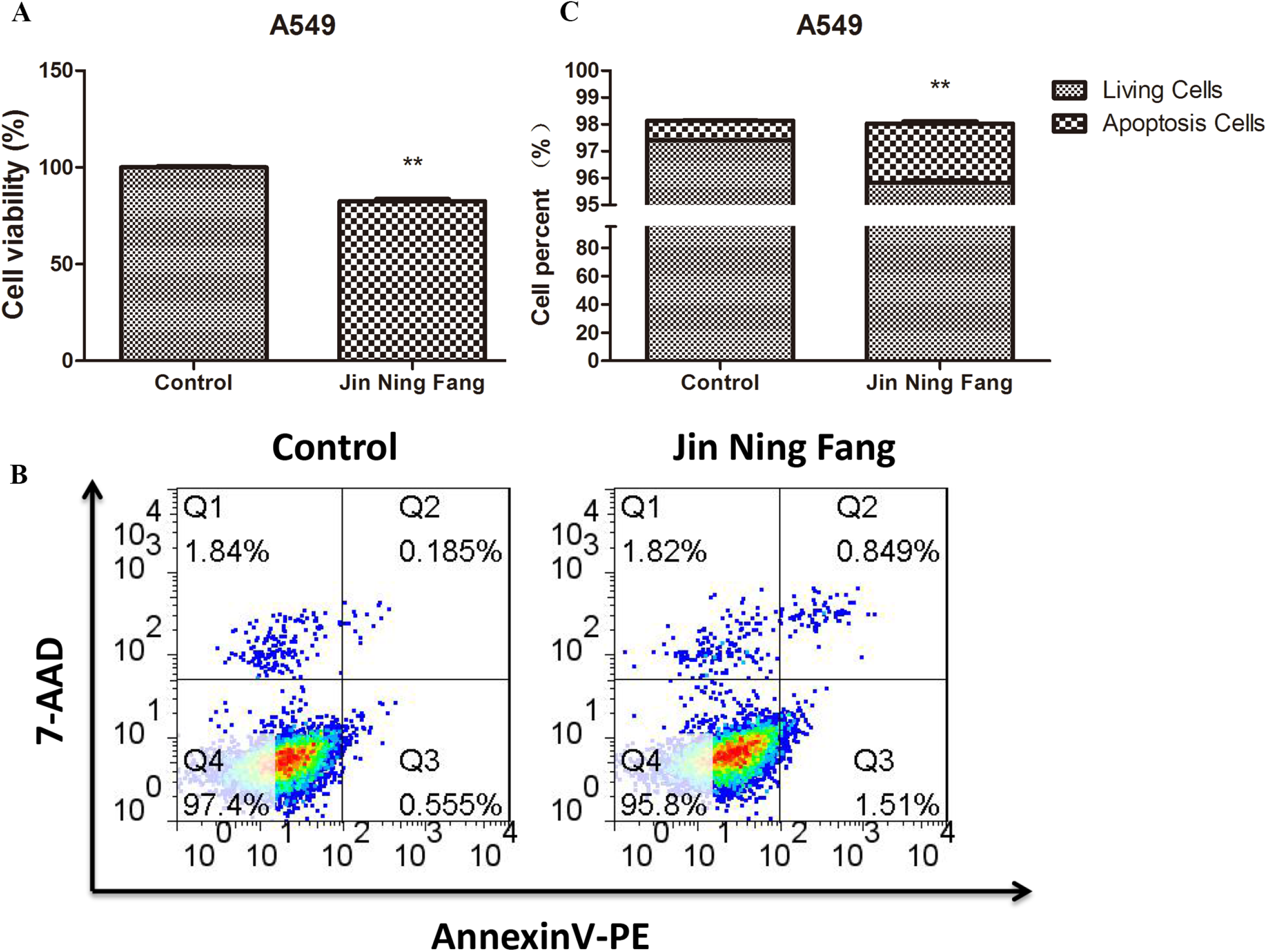
Effect of JNF on the proliferation and apoptosis of lung cancer cells. A: CCK-8 assay was performed to observe cell viability. B: Flow cytometry was used to detect the apoptosis of A549 cells. C: Quantification of living cells and apoptosis cells. Data were represented as mean ± SD. ** P < 0.01.

### Effect of JNF on several genes mRNA expression level in lung cancer cells

In order to observe the effect of JNF on gene expression level in lung cancer cells, several identified genes, including FDPS, PIM1, VCAM1, SLC29A1, NQO1, ANPEP, ESR1, and AR, were detected by qPCR. Results showed that JNF could significantly decrease the expression level of FDPS, PIM1, VCAM1, SLC29A1, NQO1, and ESR1 (all P < 0.05), while JNF markedly increase the mRNA level of AR and ANPEP (all P < 0.05, Figure 8).

**Figure 8.**
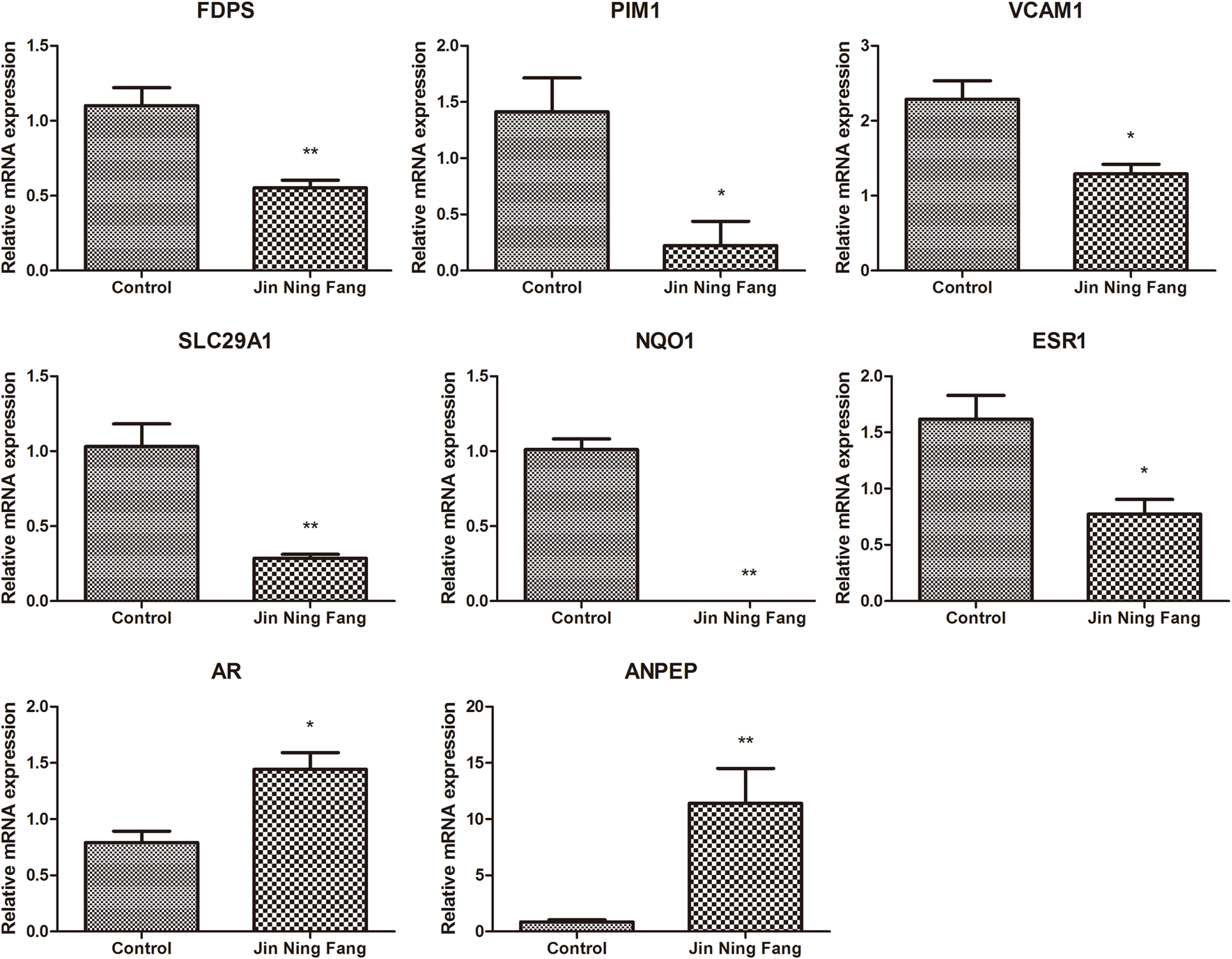
The relative mRNA expression of several genes was determined by qPCR. Data were represented as mean ± SD. * P < 0.05, ** P < 0.01.

## Discussion

In our study, two key ingredients of JNF, namely *Ostrea gigas* Thunberg (oyster) and *Salvia chinensis* Benth. (Chinese sage herb), were found to play an important role in the decoction’s therapeutic effect against lung cancer. By overlapping the target genes of the compounds in JNF and the genes related to lung cancer, 25 candidate genes were obtained. PPI analysis revealed genes such as AKT1, AR, and ESR1 as hub genes. In addition, the pharmacology network revealed that quercetin, calcium carbonate, and beta-sitosterol might be the key compounds in JNF. Furthermore, molecular docking analysis indicated that beta-sitosterol docked well with the three key targets, namely AKT1, AR, and ESR1. Moreover, we also observed that JNF could inhibit proliferation and promotes apoptosis of lung cancer cells, which further confirmed the anti-cancer effect of JNF. Importantly, the expression of several genes, including FDPS, PIM1, VCAM1, SLC29A1, NQO1, ANPEP, ESR1, and AR, was significantly altered after JNF treatment.

Results showed that quercetin, calcium carbonate, and beta-sitosterol might be considered as the key compounds in the anti-lung cancer effect of JNF. Quercetin is a flavanol and well known to be closely associated with proteins involved in apoptosis, cell cycle, detoxification, antioxidant replication, and angiogenesis (Kashyap et al. 2016). Chang *et al*. showed that quercetin, by inhibiting Akt activation, suppresses the metastatic ability of lung cancer (Chang et al. 2017). In addition, beta-sitosterol is widely known as a medicine derived from a traditional plant. Previous studies showed that beta-sitosterol regulates various pathways related to cell signaling, such as cell cycle, cell apoptosis, and cell proliferation (Ulbricht 2016). This compound also has anticancer properties against multiple cancers, including breast and lung cancers (Bin Sayeed and Ameen 2015). Moreover, in a previous study, calcium carbonate was used as a drug carrier targeting cancer cells (Maleki Dizaj et al. 2019). Taken together, the above studies further proved the potential role of these compounds in lung cancer treatment.

Chinese sage and oyster were the two important ingredients discovered in the pharmacological network that attracted our attention. Chinese sage herb has been widely used in NSCLC treatment (Maleki Dizaj et al. 2019). For example, Xu *et al*. showed that Chinese sage herb combined with chemotherapy is favorable for improving the life quality and prolonging the survival time of patients with advanced NSCLC (Xu et al. 2011). The other ingredient, oyster, which is a good source of nutrient, has also been shown to treat various cancers, including oral (Chen et al. 2016) and liver (Bin Sayeed and Ameen 2015) cancers. Furthermore, Gastineau *et al*. showed that the active compounds of oyster are effective in slowing or inhibiting the proliferation of cancer cells (Gastineau et al. 2012). This finding might provide some theoretical guidance for the development of novel drugs.

In the present study, we also identified several hub genes from both the PPI network and pharmacological network: AKT1, AR, ESR1, FDPS, and VCAM1. Liu *et al*. showed that AKT1 amplification and the mammalian target of the rapamycin (mTOR) signaling pathway play an important role in the acquirement of cis-diaminodichloroplatinum (II) (CDDP) resistance in human lung cancer cells, thereby providing a potential therapeutic target for overcoming CDDP resistance (Liu et al. 2007). Moreover, a study showed that AKT1 plays important regulatory roles in cell growth and migration, which have been recognized as targets of anticancer treatment in recent years (Lee et al. 2011). Notably, cell cycle, apoptosis, and angiogenesis pathways are important prognostic markers in the early stage of NSCLC (Singhal et al. 2005). Consistent with previous studies, our present study also showed that the target genes, such as AKT1, were significantly enriched in the platinum drug resistance and apoptosis pathways. Thus, we hypothesized that AKT1 might play a role in the anti-lung cancer of JNF by altering the platinum drug resistance and apoptosis pathways. AR is mainly associated with the regulation of male sex steroids, and its expression is recognized to be responsible for sex differences in lung development (Mikkonen et al. 2010). Ki67 is an important prognostic factor for predicting the survival of patients with lung cancer (Ciancio et al. 2012b). In lung cancer, although AR expression is not independently associated with outcome, the expression status of the gene might be related to the effect of Ki67 on the outcome (Ciancio et al. 2012a). Similarly, the methylation ratio of ESR1, a tumor suppressor gene, is related to the prognosis of lung cancer patients (Suga et al. 2008). Lin and his colleagues revealed that ESR1 methylation occurred more frequently in stage I NSCLC tissue than in non-cancerous lesions (Lin et al. 2009). Our data showed that ESR1 was enriched in the estrogen signaling pathway, and data from previous research showed that estrogens and growth factors promote lung cancer progression (Márquez-Garbán et al. 2007). FDPS (also called FPPS) is a key enzyme in the mevalonate pathway, which has been recently reported to serve a role in cancer progression. Lin et al. (Lin et al. 2018) revealed that the expression level of FPPS was significantly correlated with the TNM staging and metastasis of NSCLC, and FPPS mediated transforming growth factor β-induced NSCLC cell invasion and epithelial-to-mesenchymal transition process. Moreover, VCAM1 is a member of the Ig superfamily and encodes a cell surface sialoglycoprotein expressed by cytokine-activated endothelium. Previous study indicated that VCAM1 was overexpressed in the lung cancer-associated fibroblasts; in addition, VCAM1 could promote the growth and invasion of lung cancer cells via activating the AKT and MAPK signaling (Zhou et al. 2020). In this study, we observed that the level of VCAM1 was significantly decreased in the JNF-treated lung cancer cell, which further suggested that VCAM1 might be a therapeutic target for lung cancer. Taken together, these studies highlighted the direct and indirect relationship between the screened genes and treatment of lung cancer, indicating that these genes might be potential targets of JNF for the treatment of lung cancer.

Molecular docking is the most widely used method for calculating protein-compound interactions, and we performed this assay using Autodock to explore probable binding modes. Our results indicated that beta-sitosterol had a high binding affinity for AKT1, AR, and ESR1. These findings also indirectly showed that the results of molecular docking were consistent with those of network pharmacology, which further verified the reliability of the targets predicted via network pharmacology.

This is the first time to study the potential pharmacological mechanisms of JNF in treating lung cancer. We not only screened the potential pharmacological targets of JNF using bioinformatics analyses, but also validated the results at the cellular level. However, our study has some limitations. First, lung cancer includes many subtypes, but we have not conducted detailed studies on the different types. Second, the identified targets of JNF is involved in some signaling pathways, such as prolactin, neurotrophin signaling pathways, while the specific regulatory mechanism has not been studied in detail. Finally, our findings have not been validated *in vivo* experiments. Therefore, the focus of our follow-up research is to solve these limitations.

## Conclusion

In conclusion, the anti-lung cancer mechanisms of JNF decoction was explored via network pharmacological analysis. The results showed that quercetin, calcium carbonate, and beta-sitosterol were key compounds in the decoction. The hub genes, including AKT1, AR, ESR1, FDPS, and VCAM1, were target genes associated with the anti-lung cancer effect of JNF decoction. These findings will provide a theoretical basis for the pharmacological effects of JNFJNFJNF against lung cancer.

## Ethics Approval

All animal experiments were performed in accordance with the NIH Guide for the Care and Use of Laboratory Animals and the procedures were approved by the animal ethic committee of Longhua Hospital Affiliated to Shanghai TCM University.

## Informed consent

Not applicable.

## Data Availability Statement

All data generated or analysed during this study are included in this published article.

## Authors’ contributions

Conception and design of the research: Chunxiao Wu, Qi Bao

Acquisition of data: Qiquan Yu

Analysis and interpretation of data: Weizhen Shou, Wentao Guo

Statistical analysis: Kun Zhang, Yang Li

Drafting the manuscript: Chunxiao Wu

Revision of manuscript for important intellectual content: Qi Bao

## Supplementary materials

**Table S1.**
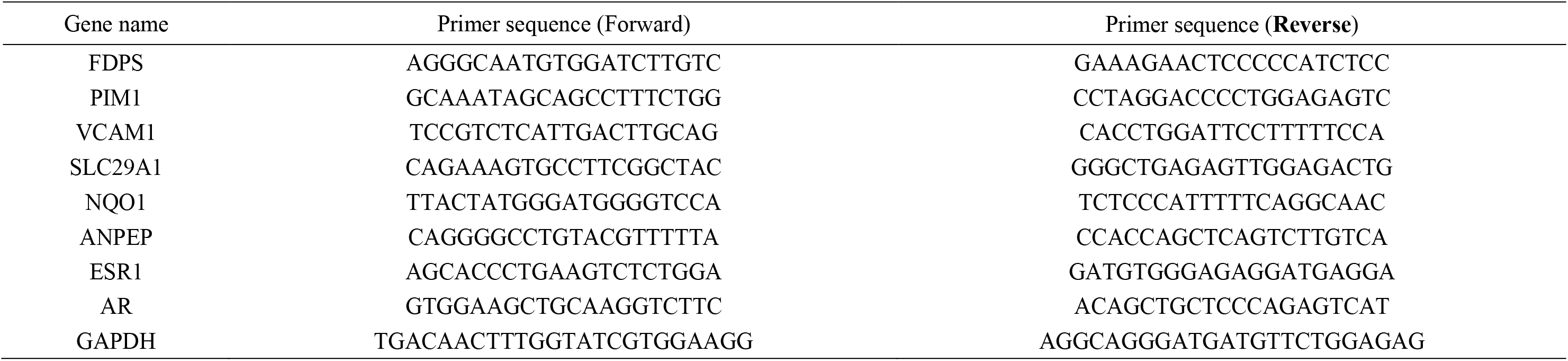
Specific primer sequences for real-time PCR

**Table S2.**
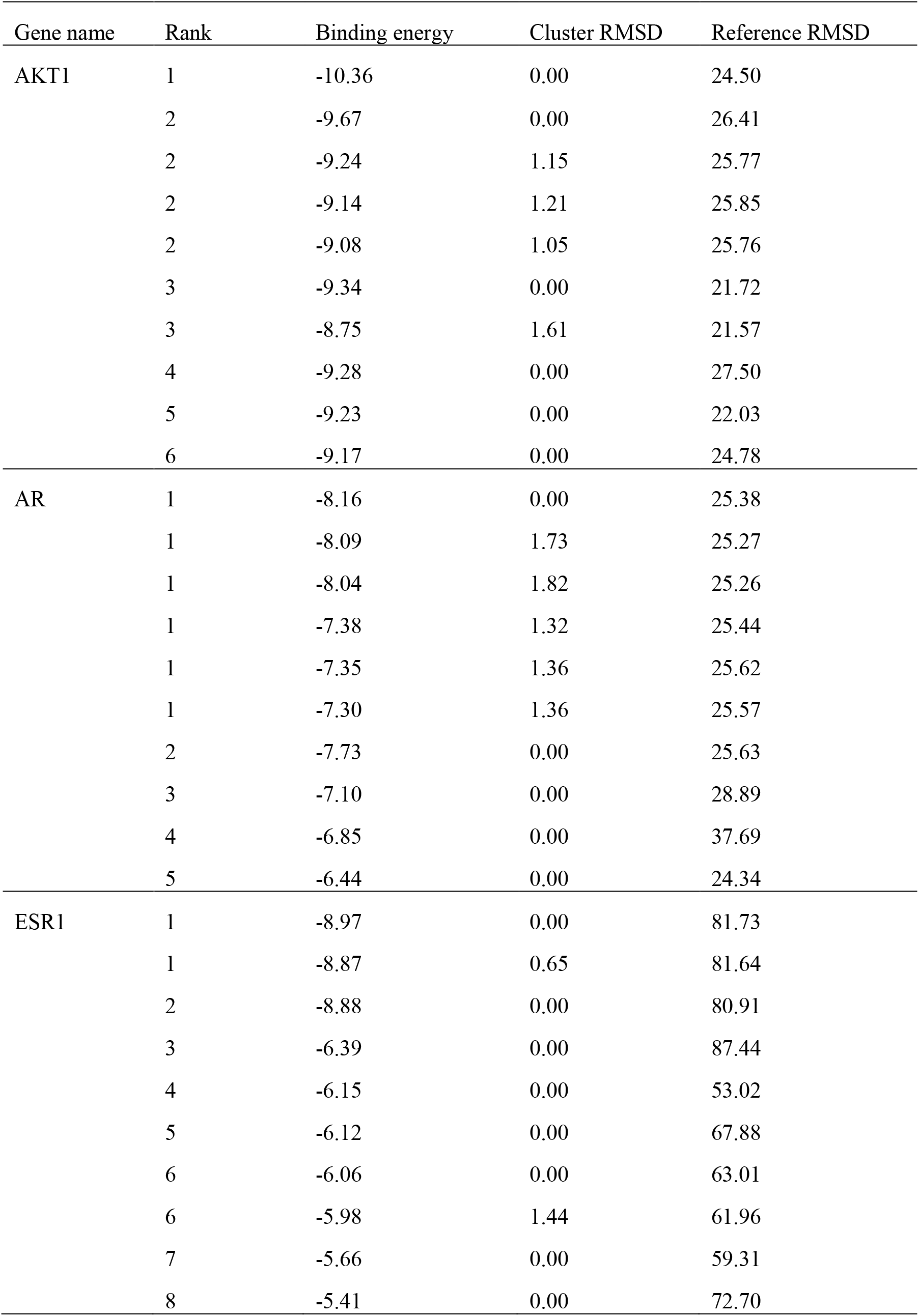
The detail score of molecular docking models

| Gene name | Rank | Binding energy | Cluster RMSD | Reference RMSD |
| --- | --- | --- | --- | --- |
| AKT1 | 1 | -10.36 | 0.00 | 24.50 |
|  | 2 | -9.67 | 0.00 | 26.41 |
|  | 2 | -9.24 | 1.15 | 25.77 |
|  | 2 | -9.14 | 1.21 | 25.85 |
|  | 2 | -9.08 | 1.05 | 25.76 |
|  | 3 | -9.34 | 0.00 | 21.72 |
|  | 3 | -8.75 | 1.61 | 21.57 |
|  | 4 | -9.28 | 0.00 | 27.50 |
|  | 5 | -9.23 | 0.00 | 22.03 |
|  | 6 | -9.17 | 0.00 | 24.78 |
| AR | 1 | -8.16 | 0.00 | 25.38 |
|  | 1 | -8.09 | 1.73 | 25.27 |
|  | 1 | -8.04 | 1.82 | 25.26 |
|  | 1 | -7.38 | 1.32 | 25.44 |
|  | 1 | -7.35 | 1.36 | 25.62 |
|  | 1 | -7.30 | 1.36 | 25.57 |
|  | 2 | -7.73 | 0.00 | 25.63 |
|  | 3 | -7.10 | 0.00 | 28.89 |
|  | 4 | -6.85 | 0.00 | 37.69 |
|  | 5 | -6.44 | 0.00 | 24.34 |
| ESR1 | 1 | -8.97 | 0.00 | 81.73 |
|  | 1 | -8.87 | 0.65 | 81.64 |
|  | 2 | -8.88 | 0.00 | 80.91 |
|  | 3 | -6.39 | 0.00 | 87.44 |
|  | 4 | -6.15 | 0.00 | 53.02 |
|  | 5 | -6.12 | 0.00 | 67.88 |
|  | 6 | -6.06 | 0.00 | 63.01 |
|  | 6 | -5.98 | 1.44 | 61.96 |
|  | 7 | -5.66 | 0.00 | 59.31 |
|  | 8 | -5.41 | 0.00 | 72.70 |

